# How tool combinations in different pipeline versions affect the outcome in RNA-seq analysis

**DOI:** 10.1101/2023.10.04.560168

**Authors:** Louisa Wessels Perelo, Gisela Gabernet, Daniel Straub, Sven Nahnsen

## Abstract

Data analysis tools are continuously changed and improved over time. In order to test how these changes influence the comparability between analyses, the output of different workflow options of the nf-core/rnaseq pipeline were compared. Five different pipeline settings (STAR+Salmon, STAR+RSEM, STAR+featureCounts, HiSAT+featureCounts, pseudoaligner Salmon) were run on three datasets (human, Arabidopsis, zebrafish) containing spike-ins of the External RNA Control Consortium (ERCC). Fold change ratios and differential expression of genes and spike-ins were used for comparative analyses of the different tools and versions settings of the pipeline. An overlap of 85% for differential gene classification between pipelines could be shown. Genes interpreted with a bias were mostly those present at lower concentration. Also, the number of isoforms and exons per gene were determinants. Previous pipeline versions using featureCounts showed a higher sensitivity to detect one-isoform genes like ERCC. To ensure data comparability in long-term analysis series it would be recommendable to either stay with the pipeline version the series was initialized with or to run both versions during a transition time in order to ensure that the target genes are addressed the same way.

## INTRODUCTION

The nf-core framework offers standardized, portable and reproducible pipelines for bioinformatic analysis workflows (1). The nf-core/rnaseq pipeline (2) is one of the most utilized pipelines in the nf-core portfolio, according to the number GitHub contributors, stargazers and users associated to its Slack channel. Its purpose is to analyse RNA sequencing data from any organism for which a reference genome and annotation is available. The pipeline takes care about data pre-processing as well as quality control and offers multiple alignment and quantifications routes. It does not include steps to analyse differential gene expression.

Several tools and workflows are available to perform the three main steps in RNA-seq analysis: (i) align reads to the reference genome, (ii) quantify genes and transcripts and (iii) identify differentially expressed genes. Some of the most popular tools used today, like STAR (3), Salmon (4) or featureCounts (5), have been released between 2013 and 2015. Benchmarking studies have been published in the years following, attributing influence on the analysis output to the tools of each of the three steps, although to different degrees, depending on the study. Williams et al. (6) tested nine aligner tools, twelve expression modelers and thirteen tools for the detection of differential expression in 219 combinatorial implementations. They found a significant heterogeneity in the outcome of these workflows and attributed this mainly to differential gene expression identification and less to the choice of read aligner and expression modeler. Also, Teng et al. (7) confirmed that mapping algorithms had a comparatively small effect on differential gene expression discovery. Other studies showed that these two latter steps are equally important for the outcome of differential gene expression analysis. Srivastava et al. (8) showed the influence of mapping and alignment approaches on the variation of quantification in experimental datasets, which in turn is crucial for differential gene expression estimation. The RGASP consortium (9) and Baruzzo et al. (10) evaluated tools for the alignment step. Both found major performance differences between the aligners evaluated. Alignment outcome varied with genome complexity and choice of parameter settings, where default values of the tools were not always set to an optimal choice. Aligners based on tools that were designed for DNA and do not consider intron-sized gaps, varied most in performance. They recommended that benchmarking of tools and pipelines should be updated regularly, due to the fast development in the field. In 2021, Sarantopoulou et al. (11) published an extensive benchmark of quantification tools, comparing six most popular isoform quantification methods. Tools performance was highly dependent on the dataset structure. They identified length and sequence compression complexity as parameters with the greatest impact on quantification accuracy.

The nf-core pipelines are constantly reviewed and updated by the open-source nf-core community. The nf-core/rnaseq pipeline was first released in 2018 and is currently available at version 3.x (latest 3.12.0, in July 2023). All versions are available on the pipelines GitHub repository (https://github.com/nf-core/rnaseq) and can be run by specifying the pipeline version in the run command (e.g. `-r 1.4.2`). According to the discussions in the developer community of the nf-core/rnaseq pipeline (Slack), the version v3.x pipelines did not implement featureCounts anymore, because the tool presents disadvantages for organisms where gene splicing occurs (11, 12). Therefore, it was deprecated and replaced by approaches that use statistical modules, like Salmon and RSEM. In the version 3.x releases, the user can choose between the aligners HiSAT2 (13) without quantification, STAR (3) in combination with quantification by Salmon (4) or RSEM (14), as well as the pseudo-alignment and quantification by Salmon, while in the previous version v1.x releases, the aligner STAR and HiSAT are available with quantification by either featureCounts, StringTie (15), Salmon or tximport (16).

Baruzzo et al. (10) recommended to monitor pipeline performance following tool updates and changes, because such a comparison is necessary for the pipeline users to interpret data generated with different versions. Therefore, the present analysis has the objective to investigate the influence of the different tools on the analysis result when migrating from v1.x to v3.x releases of the nf-core/rnaseq pipeline. Five different pipeline settings were run on three publicly available datasets from different organisms (human, plant, fish) of varying sizes (191.41 Gigabases (Gb), 119.55 Gb, 27.46 Gb) containing spike-ins of the External RNA Control Consortium (ERCC) (17). Fold change ratios and differential expression of genes and spike-ins were determined with DESeq2 (18) and used for comparative analyses of the different tools and versions settings of the nf-core/rnaseq pipeline.

## MATERIAL AND METHODS

### Datasets

Three public RNA-seq datasets from different organisms, containing External RNA Control Consortium (ERCC) spike-ins, were used in the analysis. One dataset was derived from human cell lines, one from *Arabidopsis thaliana* seedlings and the third from zebrafish. The ERCC spike-in control is a mixture of 92 synthetic polyadenylated (polyA) oligonucleotides of 250-2000 nucleotides, that are meant to resemble human transcripts (19). There are two mixtures (mix 1 and mix 2) with differing molar concentrations of these sequences. The concentrations of the transcripts in each spike-in mix span an approximately 106-fold concentration range (0.0143 to 30000 attomoles/μl) at defined molar concentration ratios between the two mixtures, distributed into 4 subgroups (4.0, 1.0, 0.67 and 0.5). Reference fasta and gtf files of the three organisms were prepared by adding the ERCC sequences and annotations (20).

#### Human cell dataset

The human cell dataset consists of the SEQC benchmark data set and data from the ENCODE project (21). The Sequencing Quality Control (SEQC) consortium generated two datasets (Group A and Group B) from two reference RNA samples in order to evaluate transcriptome profiling by next-generation sequencing technology. Each sample contains one of the reference RNA sources and a set of synthetic ERCC RNAs at known concentrations. The ‘Group A’ dataset contains 5 replicates of the Strategene Universal Human Reference RNA (UHRR), which is composed of total RNA from 10 human cell lines, with 2% by volume of ERCC mix 1. The ‘Group B’ dataset includes 5 replicate samples of the Ambion Human Brain Reference RNA (HBRR) with 2% by volume of ERCC mix 2.

#### Arabidopsis dataset

The Arabidopsis dataset was generated by Califar et al. (22) and compared the transcriptome of Arabidopsis embryos (cultivar: Wassilewskija, WS) during orbital flights to ground controls. ERCC mix 1 was added to flight samples and ERCC mix 2 to ground control samples. The controls were added with a concentration of 1 μl of 1:2000 diluted RNA spike-in ERCC spike to 50 ng of total RNA, which is half the amount suggested in the ERCC user guide.

#### Danio rerio dataset

The zebrafish dataset was published by Schall et al. (23). In this study a zebrafish model for Short Bowel Syndrome (SBS) was used to test the hypothesis that acute SBS has significant effects on gene expression associated with proliferation, inflammation, bile acid synthesis and immune system activation. The zebrafish were grouped into either SBS surgery (n□=□29, ERCC mix 2) or sham surgery (n□=□28, ERCC mix 1) groups. The data used for this benchmark originated from 3 fish from each group which were harvested for RNA sequencing at 2 weeks after surgery.

#### Reference genomes and annotations

The iGenomes Ensembl references for *Homo sapiens* (GRCh37), *Arabidopsis thaliana* (TAIR10) and *Danio rerio* (GRCz10) were used for analysis after adding the ERCC information to the fasta and gtf files. Gene length, exon number and number of isoforms per gene were retrieved from the gtf files using gtftools.py (24). For gene length, the tool provides four different types of gene lengths (the mean, median and max of lengths of isoforms of a gene, and the length of merged exons of isoforms of a gene), of which the mean length was used here for comparisons. Exons are calculated for each gene by merging exons of all splice isoforms from the same gene.

### Pipeline settings: nf-core/rnaseq

Versions 1.4.2 and 3.2 of the nf-core/rnaseq pipeline were run on the institute’s computing cluster with nextflow (v22.10.4) (25) and singularity (v. 3.8.7) (26) in five settings, differing in aligner and quantification tools. For pipeline version v1.4.2 the options for alignment by STAR or Hisat2 were used, with quantification by featureCounts. For pipeline version v3.2 alignment was done by STAR and quantification by either Salmon or RSEM. The fifth setting executed employed the pseudo aligner option using Salmon only (see full commands in Supplementary Table S1).

### Downstream analysis

#### Differential expression

The qbic-pipelines/rnadeseq pipeline (v2.0.1, https://github.com/qbic-pipelines/rnadeseq) was used to apply downstream analysis with DESeq2 (v1.34.0) to identify differentially expressed (DE) genes (see full commands in Supplementary Table S1). Resulting log2 featurecounts and gene expression classification, as well as baseMean values were used for comparison statistics.

#### Accuracy and sensitivity of pipeline performance

Comparing the known log2 ERCC spike-in concentration with the measured log2 Transcripts Per Million (TPM) can be used as an indicator for measurement accuracy and sensitivity (17, 19). Accuracy is determined by the slope of the linear regression between these two parameters. The closer the regression slope is to 1, the more accurately the pipeline predicted the relative abundance of the ERCC spike-ins. Sensitivity of the analysis workflow can be estimated by determination of the lowest limit of detection (LLD). It is defined as the molar amount of ERCC transcripts detected in a sample above a certain threshold value. When defining a sensitivity threshold of 1 TPM (log2 TPM=0), the LLD corresponds to the Y-axis value, where the regression line crosses the X-axis. This results in the log2 number of control molecules detected per concentration of Poly(A)RNA (ERCC spike-in) added to the sample (19, 27). The higher the LLD, the more molecules were detected based on the same spike-in concentration in the sample, thus being an indicator of higher analytic sensitivity.

#### Computation efficiency

All pipeline runs were executed on the institute’s computing cluster, which consists of 28 nodes in total (24 regular 32Cores/64Threads and 4 HighMem 64Cores/128Threads) and a Parallel BeeGFS Filesystem with a total capacity of 400TB. CPU hours and total memory usage were retrieved from the files in the pipelines output folder </pipeline_info/>, where the runs’ CPU hours can be found in the <execution_report.html> and total memory can be calculated from the information given in the <execution_trace.txt> file produced by each pipeline run.

#### Statistical analysis

Analysis and visualization of the DESeq2 output was performed in a Python Jupyter Notebook (6.3.0)(28), applying the packages pandas (1.2.4), numpy (1.20.2), scipy.stats (1.7.0) and scikit-learn (1.0). Graphs were generated with the python packages matplotlib (3.3.4) and seaborn (0.11.2). Venn diagrams were drawn using the R (4.2.2) library VennDiagram (1.7.3).

Gene classification as differentially expressed (DE) or not (not_DE), related gene length, baseMean and number of exons were compared on the level of the whole dataset and for ERCC spike-ins only. The DESeq parameter baseMean was used as an estimate of the dispersion of a gene, as an estimation of sequencing depth. The baseMean is defined as the ‘average of the normalized count values, divided by size factors, taken over all samples’ (29).

One-way ANOVA was applied to compare the log2 feature count values of all pipeline outcomes. In case of significant differences, the Tukey-Kramer post hoc test was applied for pairwise comparisons in order to determine which pipeline settings were differing from each other. Gene length, exon number and number of isoforms were compared as possible causes of pipeline differences by applying the independent t-test. The aim was to investigate if these parameters correlated with the classification of genes as DE or not_DE. It was also tested if these parameters were associated with the concordant classification of genes by all pipelines (concordant vs non-concordant).

ERCC sequence recovery was determined and ERCC estimates were used to calculate quality indicators like the root mean squared error between expected and observed log2 fold-change, as well as the lower limit of detection (LLD) (17, 19).

## RESULTS AND DISCUSSION

### Identification of differentially expressed genes in whole datasets

#### Concordance between pipeline outputs and the influence of dataset structure

In a first step, the classification of all genes in the datasets as differentially (DE) and non-differentially expressed (not_DE) was compared for the five pipeline settings. As already observed in previous benchmarks studies (6, 8–11), log2 feature count values and hence the number of differentially expressed genes varied between pipelines and tools (Fig. 1). Dataset related parameters that may influence the analysis of RNA expression include gene length, number of exons and gene isoforms, as well as sequencing depth. Short gene length, a high number of exons and a low sequencing depth were characteristic for the genes that were ambiguously classified in a benchmark study of RNA-sequencing analysis workflows (30). Also, the number of isoforms and the expression pattern of those were shown to have an influence on tool performance, especially if the short isoforms are the dominant transcripts (12).

**Figure 1.**
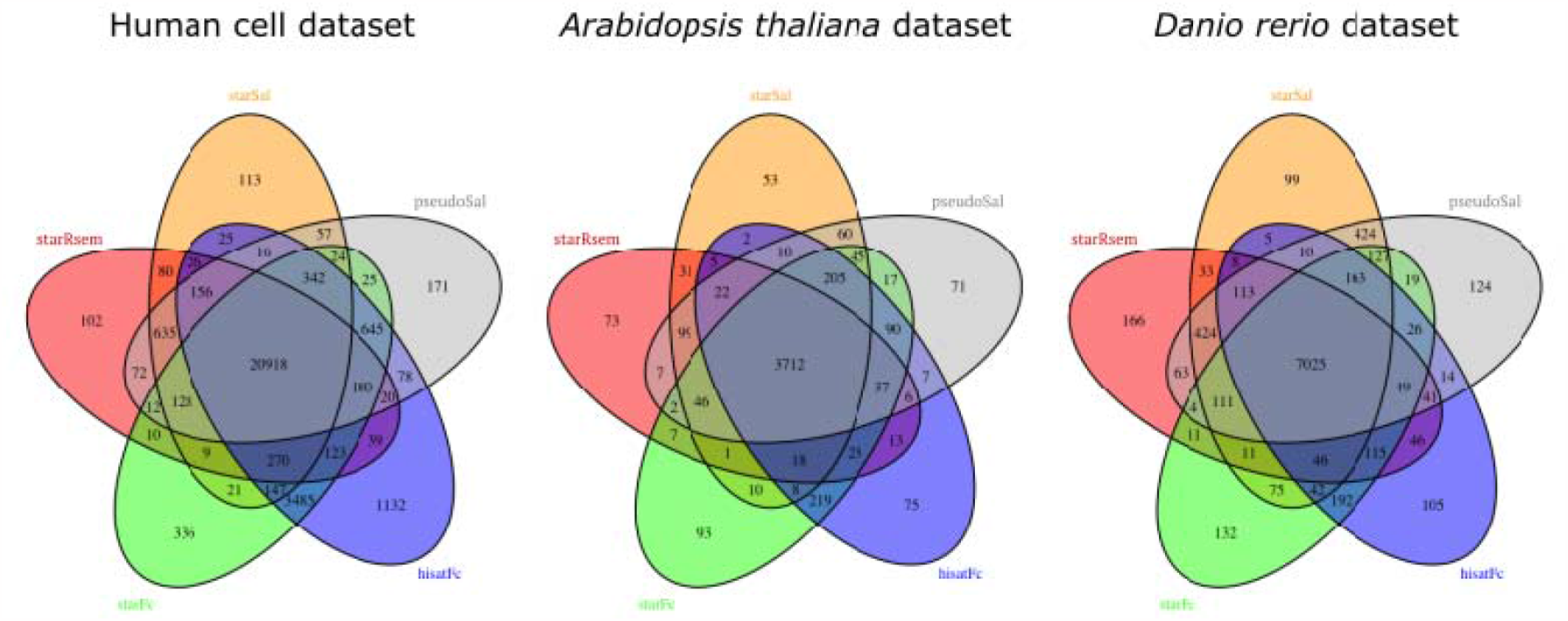
Venn diagrams showing the number of differentially expressed (DE) genes identified by pipeline for each dataset. The number in the center represents all genes identified as differentially expressed by all pipelines. All other overlaps include the number of genes identified as DE by the pipelines represented in the overlap.

The counts done by gtftools.py (24) for the organisms’ reference genomes used in this study were differing in size, number of exons and isoforms per gene. These counts from the gtf-files include not only protein-coding genes, but also the non-coding genes in the reference genome, including non-protein coding RNA-producing genes. The human reference genome was the largest (∼58000 genes) and had an elevated number of genes with a low exon count: 40% of all genes in this dataset had one exon, 71% up to five and a maximum of 367 exons were counted in one gene (Supplementary Fig. S1). Regarding the number of isoforms for one gene, the human reference genome stood out for having more genes with a high number of isoforms, with a maximum of 82 transcripts for one gene (Supplementary Fig. S2). The Arabidopsis genome was of medium size (∼34000 genes) with a comparable distribution of exons (34% containing one, 73% up to 5 exons), a maximum number of 77 exons in one gene and a maximum of 27 isoforms for the most varied gene. The zebrafish genome on the contrary was the smallest genome (∼29000 genes) with a higher exon number per gene: only 14% of genes contained one exon, 41% up to five exons and the maximum number within one gene was 590. The highest number of isoforms for one gene was 20.

In the human cell dataset, the largest amount of DE genes was identified, while the Arabidopsis dataset was the one with the least DE genes. It is also noteworthy that in the human cell dataset, a larger number of DE genes was detected by the version v1.4.x pipelines (star+featureCounts, hisat+featureCounts), being more pronounced after alignment with Hisat2 (Fig. 1). The use of featureCounts is supposed to have a higher performance on one-isoform genes (11), which in the human reference genome make up 62% of all genes . Not shown in the Venn Diagrams are the non-differentially expressed genes (non_DE) identified in all pipeline runs, which sum up to 18387, 19054 and 13625 for the human cell, Arabidopsis and zebrafish datasets, respectively.

The general recommendation given by authors of benchmarking studies (11, 12) is to prefer tools that use statistical modules, like Salmon and RSEM, over featureCounts, considering them more reliable. One study (12) compared featureCounts with RSEM and attributed the difference between the two approaches primarily to the number of transcripts in a gene. The more transcripts a gene had, the larger the difference tended to be, especially when the short isoforms were the dominant transcripts. The study of Sarantopoulou et al. (11) attributed accuracy differences mainly to length and sequence compression complexity and to a lesser extent to the number of isoforms. Comparing six quantification tools on idealized and realistic datasets, the tools that exhibited the highest accuracy on idealized data, did not perform dramatically better on the more realistic data. While in their introduction the authors explicitly did not recommend featureCounts, they then concluded that their results confirmed that simpler approaches such as featureCounts appeared to do better on one-isoform genes, with the final suggestion to treat those genes separately (11).

Considering the sum of genes identified concordantly as DE or not_DE by all pipelines, a mean of 86.7% was reached for all datasets, while 0.3% to 1.4% were assigned as DE by only one pipeline setting. This is in accordance with other benchmark studies, where a range of 80% to 85% of concordance between tools was determined (11, 30). In the benchmark of quantification tools performed by Sarantopoulou et al. (11), Salmon, RSEM and Cufflinks were shown to be highly concordant with the truth up to the 80th percentile, while for featureCounts this was less. Also, Everaert et al. (30) found that about 85% of the genes showed consistent results in their benchmarking study.

DE genes only detected by a single pipeline setting had in common that they presented a significantly lower baseMean value than the concordant DE genes (one-way ANOVA, p = 4x10^-70^, 3x10^-34^, and 6x10^-14^ for human, Arabidopsis and zebrafish, respectively). This should be considered, if the purpose of the RNA-sequencing is to find low expressed genes. Gene length and exon number did not show a consistent pattern of correlation with the concurring assignment of genes by tool combinations. Concordant DE genes had a greater mean length, with a significant difference in the human cell and Arabidopsis datasets (t-test p<3x10^-5^), but not in the zebrafish dataset (p=0.13). In the human cell dataset, the genes identified concordantly as DE by all pipelines also contained a significantly higher number of exons than the DE genes identified only by a single pipeline setting (t-test p=2x10^-43^), corresponding with previous benchmark finding (30). Regarding the number of isoforms for one gene, the human cell dataset showed a significantly higher number of isoforms (p=1.3x10^-68^) for the genes concordantly identified as DE by all pipeline setting. This could not be confirmed in the other two datasets, matching findings for the tools compared by Kanitz et al. (31), including RSEM, where the output was “largely not affected by structural features” as exon number or transcript length, but who also confirmed that abundant transcripts, here indicated by a higher baseMean value, were quantified more accurately compared to rare ones.

#### Log2 fold change: correlation between pipelines

In a second step, the log2 fold-change values for all genes were compared between all analysis settings. The deviation of one pipeline setting against another was determined by setting one measurement as ground truth and calculating the root mean squared error (RMSE) and the R^2^ of the second setting against it (Table 1, Supplementary Figure S3). Best correlation and lowest errors occurred between pipelines of the same version, with an average RMSE of 0.53±0.1. Increased differences in terms of a significantly higher RMSE (t-test p=5.4x10^-8^) occurred when comparing the outcome of the 1.4.2 version to 3.2 version pipelines (RMSE 0.92±0.16). This finding varied with the dataset and was least pronounced in the zebrafish dataset (Table 1). It is noteworthy that the outcome of star+salmon for alignment and quantification showed a high correlation with the results from the much faster pseudo-aligner salmon (Table 1: human cell dataset 95%, Arabidopsis 0.85%, zebrafish 0.94%). Together with the data on CPU and memory demand for the pseudoaligner (see below), this option turns into an interesting option for high throughput data.

**Table 1.** Root mean squared error and correlation coefficient R2 (in brackets) of observed log2 fold-change compared to a hypothetical equal outcome for two pipelines. (ssal=star+salmon, srsem=star+rsem, sfc=star+featureCounts, hfc=hisat+featureCounts, psal=pseudo-aligner salmon)

| *Homo sapiens* |  |  |  |  | *Arabidopsis thaliana* |  |  |  |  | *Danio rerio* |  |  |  |  |
| --- | --- | --- | --- | --- | --- | --- | --- | --- | --- | --- | --- | --- | --- | --- |
|  | srsem 3.2 | sfc 1.4.2. | hfc 1.4.2 | psal 3.2 |  | srsem 3.2 | sfc 1.4.2. | hfc 1.4.2 | psal 3.2 |  | srsem 3.2 | sfc 1.4.2. | hfc 1.4.2 | psal 3.2 |
| ssal 3.2 | 0.398<br>(0.97) | 0.985<br>(0.85) | 1.140<br>(0.81) | 0.545<br>(0.95) | ssal 3.2 | 0.517<br>(0.85) | 0.982<br>(0.55) | 1.010<br>(0.53) | 0.521<br>(0.85) | ssal 3.2 | 0.683<br>(0.84) | 0.854<br>(0.85) | 0.659<br>(0.84) | 0.388<br>(0.94) |
|  | srsem 3.2 | 1.010<br>(0.84) | 1.150<br>(0.81) | 0.565<br>(0.94) |  | srsem 3.2 | 1.040<br>(0.50) | 1.030<br>(0.52) | 0.675<br>(0.75) |  | srsem 3.2 | 0.755<br>(0.81) | 0.703<br>(0.83) | 0.701<br>(0.83) |
|  |  | sfc 1.4.2 | 0.486<br>(0.97) | 0.945<br>(0.86) |  |  | sfc 1.4.2 | 0.397<br>(0.92) | 0.931<br>(0.59) |  |  | sfc 1.4.2 | 0.432<br>(0.93) | 0.684<br>(0.83) |
|  |  |  | hfc 1.4.2 | 1.090<br>(0.83) |  |  |  | hfc 1.4.2 | 0.955<br>(0.57) |  |  |  | hfc 1.4.2 | 0.674<br>(0.84) |

### ERCC spike-ins: detection, pipeline sensitivity and accuracy

#### Recovery of ERCC spike-ins

The ERCC spike-in mix used in all three datasets contains 92 synthetic polyadenylated (polyA) oligonucleotide sequences of differing length, concentration and mix 1 to mix 2 concentration ratio. The number of spike-ins identified in the sequenced datasets and the accuracy with which output counts are reflecting the expected concentration and ratios were analyzed.

The Venn Diagrams in Fig. 2 give an overview of the ERCC spike-ins identified by the different pipeline settings in the three datasets. The total of all 92 ERCC spike-ins were only identified once: in the human cell dataset with the hisat+featureCounts setting. In this dataset, the v1.4.2 star+featureCounts pipeline found 90 ERCCs and the v3.2 tool combinations only detected 87 spike-ins. In the Arabidopsis dataset, the v1.4.2 pipelines also detected more ERCC spike-ins, than v3.2 output, with 79 versus 77 spike-ins. The lowest number of detected ERCC spike-ins occurred in the Arabidopsis dataset, where a total of 13 spike-in sequences were missing in any of the pipeline settings, while in the zebrafish dataset only three spike-ins were not recovered. In the latter all pipelines identified 87 of the 92 spike-ins, while the settings Star+RSEM and Star+featureCounts each identified an additional ERCC gene. Spike-in concentrations in the ERCC mixes varied between 0.0143 and 30,000 attomoles/μl. The undetected spike-in sequences were the ones with the lowest concentration in the mix. In the zebrafish dataset the spike-ins with an original concentration of <0.028 attomoles/μl were not detected, while in the Arabidopsis dataset the spike-ins of <1.83 attomoles/μl were missing, but here the ERCC mix was only added at half the concentration recommended by the manufacturer.

**Figure 2.**
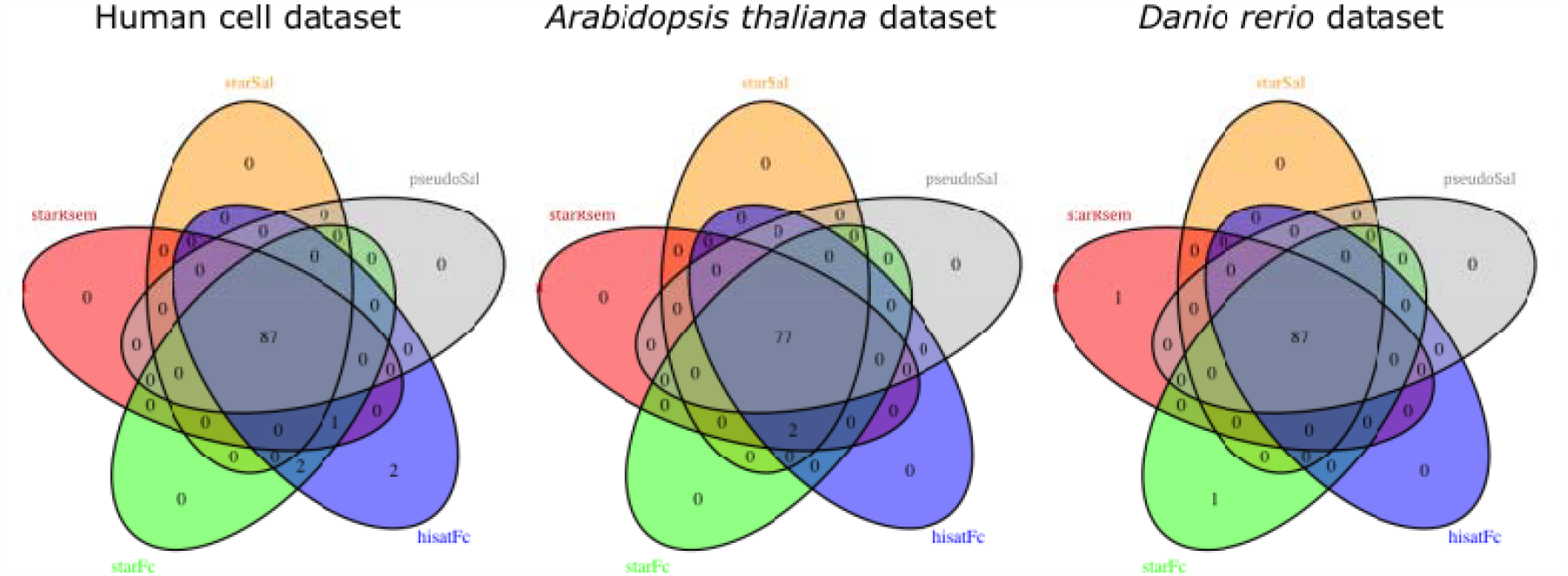
Venn diagrams showing the number of ERCC spike-ins detected by each pipeline version and setting (expected: 92). The number in the center represents spike-ins detected by all pipelines. All other fields include the number of ERCCs identified only by the pipelines represented in the overlap.es identified as DE by the pipelines represented in the overlap.

#### Pipeline accuracy and sensitivity for ERCC indicators

In order to estimate how accurately the pipelines predicted the actual ground truth given by the concentration of the artificial ERCC spike-ins and their ratio in mix 1 and mix 2, expected and observed log2 fold-change were compared.

The parameter baseMean, used as an estimate of the dispersion of a gene and therefore related to sequencing depth, has a significant impact on the result of DE analysis (30), which could also be confirmed in this study. Thus, measured ERCC spike-ins were filtered by increasing baseMean and the root mean squared error (RMSE) of the observed to expected log2 fold-change was calculated. Figure 3 shows how the RMSE decreases with increasing baseMean for the different pipeline settings in the three datasets. Numbers above the data points indicate how many ERCC spike-ins were part of the baseMean range and were included in the error calculation.

**Figure 3.**
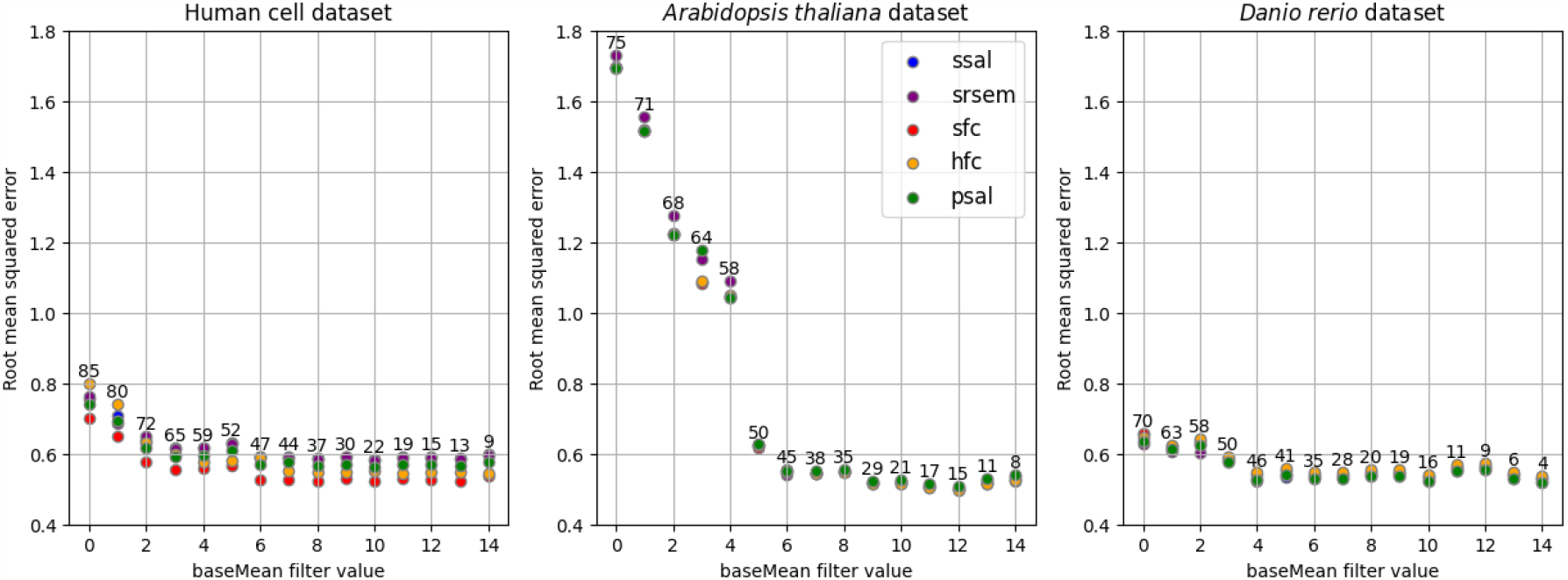
Root Mean Squared Errors (RMSE) for computed log2 fold-change of ERCC spike-ins compared to ERCC ground truth, filtering data by baseMean. The numbers above data points indicate the number of ERCCs included in the respective baseMean (>= value) range.

In the zebrafish dataset the baseMean showed little influence on the RMSE, while in the Arabidopsis dataset the error grew exponentially when including the reads with low baseMean. Filtering the data by increasing baseMean leads to an RMSE approaching a value between 0.5 and 0.6. For the human cell dataset, the error value decreased from 0.8 to below 0.6 with a baseMean equal or greater than 3, containing 65 of the 92 ERCC sequences present in the mix. In the Arabidopsis dataset a decrease from 1.7 to <0.6 was reached at a baseMean >6, including 45 ERCC sequences. Compared to other datasets, the zebrafish dataset improved less (0.7 to <0.6) with baseMean cutoff >4 with 46 ERCC sequences.

After filtering for higher baseMean values, the log2 fold-change showed a correlation to the ERCC ground truth such that the linear regression line through the data points has the same slope of 1 as the expected ground truth (Fig. 4). Without filtering this slope decreased to 0.83, 0.30 and 0.88 for the human cell, Arabidopsis and zebrafish datasets, respectively.

**Figure 4.**
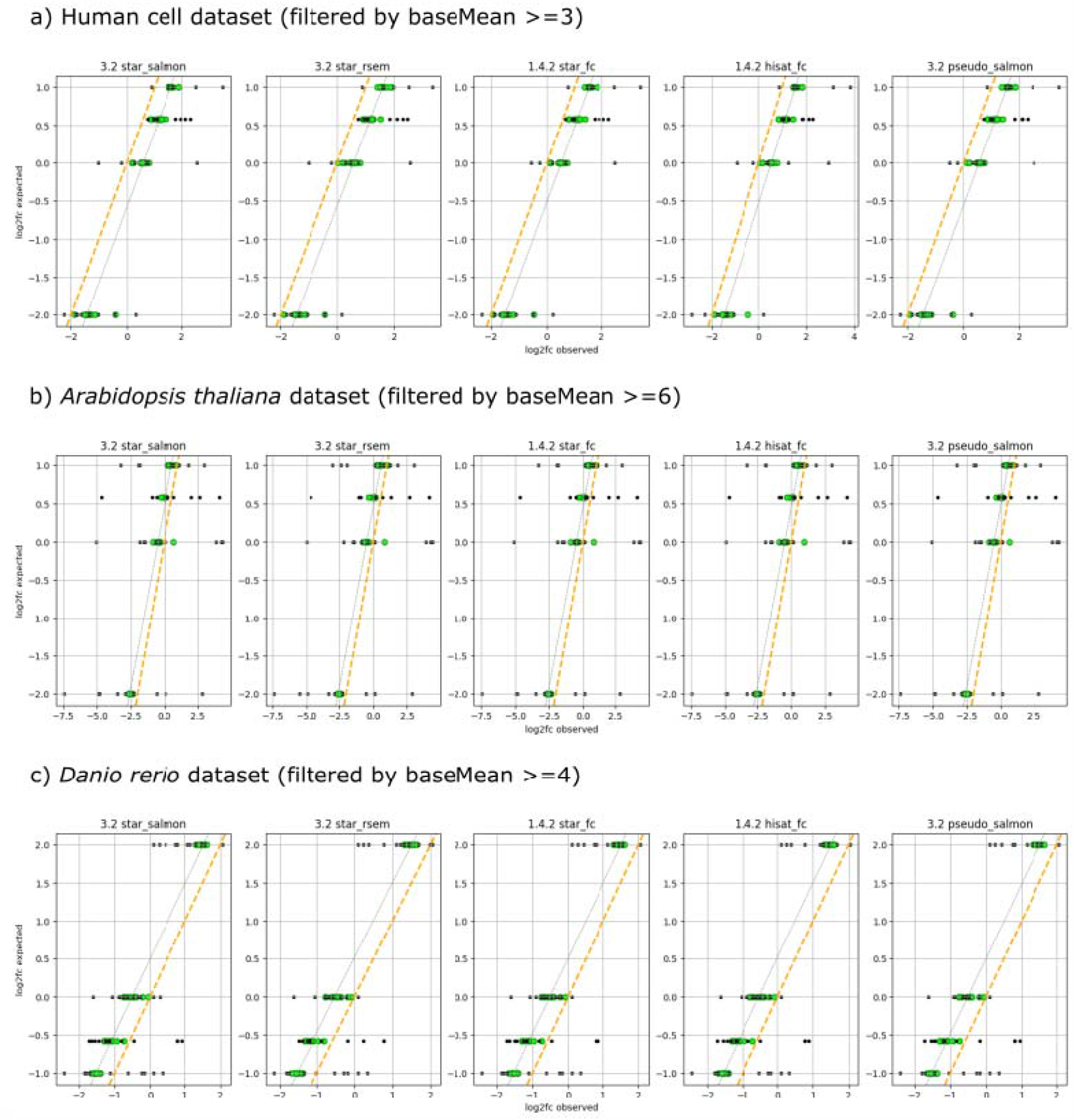
Observed and expected log2 fold-change for ERCC spike-ins, filtered by baseMean values (with root mean squared error [RMSE] to ground truth <0.6). Green: data points with a baseMean value above the cut-off, Gray: data points with a baseMean value below the cut-off. The gray dotted line indicates the linear correlation between measured log2 fold-change of the filtered (green) data points. The orange line shows the expected ground truth of the synthetic ERCC spike-ins.

Figure 4 also shows a shift of 0.5 in fold-change estimates to the left or to the right, depending on the dataset but independent of pipeline settings. The fold-changes in the human cell dataset were overestimated (regression line shifted to the right, Fig. 4a) and underestimated in the Arabidopsis (Fig. 4b) and zebrafish datasets (regression line shifted to the left, Fig. 4c). This over- or underestimation (a dataset specific systematic error) could not be explained within this benchmark setting and might be due to upstream processes like sample preparation and sequencing techniques. Further investigation with more datasets would be needed focusing on upstream sample handling procedures.

The accuracy of the measurement can be determined using the slope of a regression line between the observed log2 TPM values of ERCC spike-ins with the expected log of spike-in concentrations. The closer the regression slope is to 1, the more accurately the pipeline predicted the relative abundance of the ERCC spike-ins. The linear regression slopes for the three datasets reached values >0.9, with a minimum of 0.92 (zebrafish dataset with pseudo-aligner salmon) and a maximum of 0.97 (Arabidopsis dataset for v1.4.2 pipelines). In the human cell dataset, no difference was detected between pipeline settings (0.96 for all settings) (Table 2).

**Table 2:** Linear regression slope and Lowest Limit of Detection (LLD) for observed Transcripts per Million (TPM) and expected ERCC concentration values in the three datasets: human cell lines (Homo sapiens), Arabidopsis (Arabidopsis thaliana) and zebrafish (Danio rerio).

| *dataset* | *Homo sapiens* |  | *Arabidopsis thaliana* |  | *Danio rerio* |  |
| --- | --- | --- | --- | --- | --- | --- |
| *pipeline settings* | **slope** | **LLD** | **slope** | **LLD** | **slope** | **LLD** |
| **star+salmon** | 0.96 | 3.81 | 0.96 | 4.11 | 0.93 | 7.14 |
| **star+rsem** | 0.96 | 3.88 | 0.96 | 4.41 | 0.95 | 6.89 |
| **star+featureCounts** | 0.96 | 18.5 | 0.97 | 4.48 | 0.95 | 7.75 |
| **hisat+featureCounts** | 0.96 | 17.6 | 0.97 | 4.56 | 0.95 | 6.72 |
| **pseudoaligner salmon** | 0.96 | 3.78 | 0.96 | 4.05 | 0.92 | 6.81 |

The determination of the LLD (Table 2) gives an estimate of the method’s sensitivity. In the Arabidopsis and the zebrafish datasets determination of LLD values (Table 2) resulted in slightly higher values in the v1.4.2 settings. This difference was more pronounced in the human cell line dataset, indicating a higher sensitivity of the v1.4.2 pipelines to detect the ERCC spike-ins. This finding also corresponds to the data visualized in the Venn-Diagram (Fig. 2), where more ERCC spike-ins were detected with the v1.4.2 pipelines. The v1.4.2 pipelines use featureCounts for quantification and this tool is more sensitive to one-isoform genes (11). This is also of particular interest in prokaryote research, where gene splicing does not occur. Genes identified as differentially expressed only by the v1.4.2 pipelines in the human cell dataset had a significantly lower number of isoforms (2.2±3.6) than the genes identified as DE exclusively by v3.2 pipelines (5.3±5.8, independent t-test p=1.65x10 ^-115^).

Overall, the v1.4.2 pipeline setting using featureCounts for quantification demonstrated a higher sensitivity in detecting ERCC spike-ins. However, the ERCC spike-ins are non-spliced sequences, therefore any advantage of Salmon or RSEM regarding the differentiation of these cannot be assessed through these markers. The higher sensitivity on ERCC spike-ins is in concordance with the findings of Sarantopulou et al. (11), which confirmed a higher performance of featureCounts on one-isoform genes. The human cell dataset also is what (11) described as “idealized” data, which is expected to obtain upper bounds on the accuracy of all methods. They draw attention that in their study also brain cells were compared to liver cells, which results in an elevated amount of DE genes compared to typical experiments and therefore “may be beyond the assumptions of the DE software, particularly regarding normalization”. Therefore, the difference in tools’ performance could be more pronounced on these idealized than on more realistic data. This can also apply to the present study, where the difference in sensitivity was more pronounced in the “idealized” human cell dataset compared to the other two datasets.

#### Computation efficiency

The three datasets had very different sizes (Table 3) and thus CPU hours and memory usage differed between them. Regarding the different pipeline settings, it could be observed that the v3.2 pipelines using alignment (star+salmon and star+rsem) had the highest demand compared to the old version v1.4.2 pipelines with alignment and featureCounts for quantification. The pseudo-aligner Salmon using an estimator instead of alignment was the most efficient and fast setting, however at the cost of a slight decrease of sensitivity (Table 2). For DE gene expression the pseudo-aligner Salmon reached a correlation coefficient R2 of up to 95% when compared to the alignment-based approach with STAR and Salmon (Table 1). High correlation and computation efficiency make this option interesting for high throughput data. For small or medium sized RNA-Seq studies genome alignments are recommended (11), because then coverage plots can be examined in a genome browser.

**Table 3:** Computing statistics from pipeline runs retrieved from the nextflow tower output (CPU time and total memory).

| *Dataset*<br>(Gigabases, Gb) | **H. sapiens**<br>191.42 Gb |  | **A. thaliana**<br>119.55 Gb |  | **D. rerio**<br>27.46 Gb |  |
| --- | --- | --- | --- | --- | --- | --- |
| *pipeline settings* | CPUh | total Memory (GB) | CPUh | total Memory (GB) | CPUh | total Memory (GB) |
| **star+salmon** | 608.6 | 1379.31 | 528.7 | 412.5 | 58.3 | 779.83 |
| **star+rsem** | 930.5 | 1121.70 | 382.5 | 393.46 | 98.9 | 404.52 |
| **star+featureCounts** | 328.8 | 868.51 | 420.5 | 187.91 | 64.5 | 551.05 |
| **hisat+featureCounts** | 240.7 | 684.14 | 150.3 | 257.01 | 47.4 | 335.42 |
| **pseudoaligner salmon** | 130.6 | 257.01 | 62.0 | 26.94 | 17.0 | 147.51 |

#### Recommendations for pipeline use and development

The main motivation for this benchmark study was to find out, how the change of tools applied in a migration from the nf-core/rnaseq pipeline v1.4.2 to v3.x would influence the comparability of the output. As described above, the different tool combinations showed an overlap of around 85%, a range already reported in other studies (11, 30). Genes interpreted with a bias between pipelines were mostly those present in lower concentrations. Besides this, the overlap of the tools output was dependent on the dataset structure, were the number of transcripts per gene and the number of exons were determinants. To ensure data comparability in long-term analysis series it would be recommendable to either stay with the pipeline version the series was initialized with or to run both versions during a transition time in order to ensure that the target genes are addressed the same way. This is especially important, if the target genes are known to have a single isoform, as predominant in prokaryotes, or a low expression rate is expected.

Regarding pipeline development, it would be worth discussing to include the tool featureCounts as a quantification tool option also in the v3.x pipelines. The advantage of featureCounts on datasets with a high number of one-isoform genes has been shown here and in a previous study (11) and should also be kept in mind for the analysis of prokaryotic transcriptomes.

Running a benchmark script on a selected dataset at new pipeline releases could be used to determine how or if the pipeline performance is increased with new releases. Of the three datasets used in the present study, the human cell dataset would probably be the most informative, because here all 92 ERCC spike-ins were detected with at least one of the pipeline settings (HISAT+featureCounts). This confirms that this dataset includes reads of all spike-in sequences. In the Arabidopsis and zebrafish datasets the non-detection of ERCC sequences could mean that they are either absent in the read files due to upstream issues in sample generation and sequencing steps or that the applied pipeline tools were not able to detect them. Also, in the human cell dataset the lowest baseMean cutoff could be applied to achieve a slope close to 1 when comparing the expected and measured log2 fold-change of spike-ins (Fig. 4). As shown in this study, possible indicators of performance improvement could be the number or ERCC spike-ins recovered, slope and LLD of expected versus measured ERCC spike-ins, minimum baseMean cutoff to reach a slope close to 1 for the linear regression of observed versus expected log2 fold-change values, as well as the computation efficiency (CPU hours, total memory). The values determined in this study may serve as the baseline for future benchmark comparisons of the nf-core/rnaseq and equivalent pipelines.

## DATA AVAILABILITY

The datasets used in this study are publicly available under the following project IDs: human cell dataset (21) PRJNA214799 (GEO: GSE49712, samples SRR950078-SRR950087), *Arabidopsis thaliana dataset* (22) PRJNA674629 (GEO: GSE160846, samples SRR12980996-SRR12981001), Danio rerio dataset (23) PRJNA325275 (GEO: GSE83195, samples SRR3655791-SRR3655802).

## ACKNOWLEDGEMENTS

We thank the nf-core community in general and especially the developers and contributors to the nf-core/rnaseq pipeline for providing the platform for open-source, high-quality, reproducible bioinformatics pipelines.

