## Supplementary for "How tool combinations in different pipeline versions affect the outcome in RNA-seq analysis"

Table S1: Full commands used for the execution of the pipelines nf-core/rnaseq and qbic-pipelines/rnadeseq

| **run** | **command** |
| --- | --- |
| **https://github.com/nf-core/rnaseq** | |
| STAR+Salmon (ssal) | nextflow run nf-core/rnaseq \  -r 3.2 \  -profile singularity \  --input 'samplesheet.csv' \  --outdir 'results_ssal' \  --fasta 'genome.fa' \  --gtf 'genes.gtf' \  --aligner star_salmon |
| STAR+RSEM (srsem) | nextflow run nf-core/rnaseq \  -r 3.2 \  -profile singularity \  --input 'samplesheet.csv' \  --outdir 'results_srsem' \  --fasta 'genome.fa' \  --gtf 'genes.gtf' \  --aligner star_rsem |
| STAR + featureCounts (sfc) | nextflow run nf-core/rnaseq \  -r 1.4.2 \  -profile singularity \  --reads 'SRR*_{1,2}.fastq.gz' \  --outdir 'results_sfc' \  --fasta 'genome.fa' \  --gtf 'genes.gtf' \  --aligner salmon \ |
| HiSAT2 + featureCounts (hfc) | nextflow run nf-core/rnaseq \  -r 1.4.2 \  -profile singularity \  --reads 'SRR*_{1,2}.fastq.gz' \  --outdir 'results_hfc' \  --fasta 'genome.fa' \  --gtf 'genes.gtf' \  --aligner hisat2 |
| Pseudoaligner Salmon (psal) | nextflow run nf-core/rnaseq \  -r 3.2 \  -profile singularity \  --input 'samplesheet.csv' \  --outdir 'results_psal' \  --fasta 'genome.fa' \  --gtf 'genes.gtf' \  --pseudo_aligner salmon \  --skip_alignment true |
| **https://github.com/qbic-pipelines/rnadeseq** | |
| Quantification with featureCounts (sfc, hfc) | nextflow run qbic-pipelines/rnadeseq \  -r 2.0.1 \  -profile singularity \  --gene_counts ' results_rnaseq/featureCounts/merged_gene_counts.txt' \  --input_type 'featurecounts' \  --outdir 'results_deseq' \  --genome <Genome ID, e.g. 'GRCz10‘> \  --organism <Organism name, e.g. ‘drerio’> \  --multiqc 'results_rnaseq/multiqc.zip' \  --metadata 'samplesheet.tsv' \  --model 'model.txt' \  --project_summary 'project_summary.tsv' \  --versions 'results_rnaseq/pipeline_info/software_versions.csv' \  --skip_pathway_analysis true \  --gtf 'genes.gtf' |
| Quantification with salmon (ssal, psal) | nextflow run qbic-pipelines/rnadeseq \  -r 2.0.1 \  -profile singularity \  --gene_counts '<rnaseq_results_dir>/star_salmon’ \  --input_type 'salmon' \  --outdir 'results_deseq' \  --genome <Genome ID, e.g. 'GRCz10‘> \  --organism <Organism name, e.g. ‘drerio’> \  --multiqc '<rnaseq_results_dir>/multiqc.zip' \  --metadata 'samplesheet.tsv' \  --model 'model.txt' \  --project_summary 'project_summary.tsv' \  --versions ‘<rnaseq_results_dir>/pipeline_info/software_versions.csv' \  --skip_pathway_analysis true \  --gtf 'genes.gtf' |
| Quantification with RSEM (srsem) | nextflow run qbic-pipelines/rnadeseq \  -r 2.0.1 \  -profile singularity \  --gene_counts '<rnaseq_results_dir>/star_rsem’ \  --input_type 'rsem' \  --outdir 'results_deseq' \  --genome <Genome ID, e.g. 'GRCz10‘> \  --organism <Organism name, e.g. ‘drerio’> \  --multiqc '<rnaseq_results_dir>/multiqc.zip' \  --metadata 'samplesheet.tsv' \  --model 'model.txt' \  --project_summary 'project_summary.tsv' \  --versions ‘<rnaseq_results_dir>/pipeline_info/software_versions.csv' \  --skip_pathway_analysis true \  --gtf 'genes.gtf' |

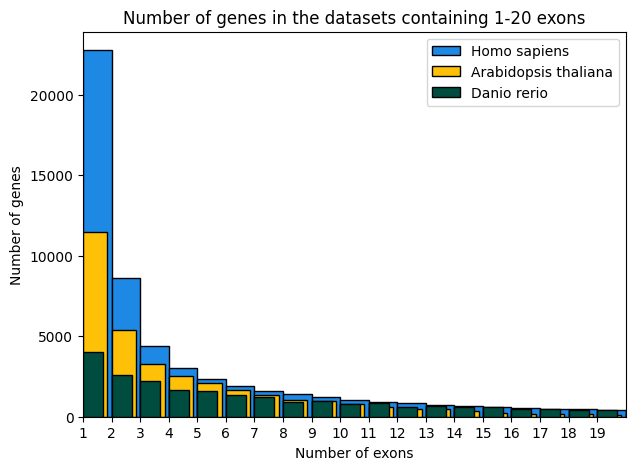

Figure S1: Number of genes containing 1-20 exons in the three datasets (Homo sapiens, Arabidopsis thaliana and Danio rerio). A maximum of 367, 77 and 590 exons in one gene were counted in the reference datasets for Homo sapiens, Arabidopsis thaliana and Danio rerio, respectively.

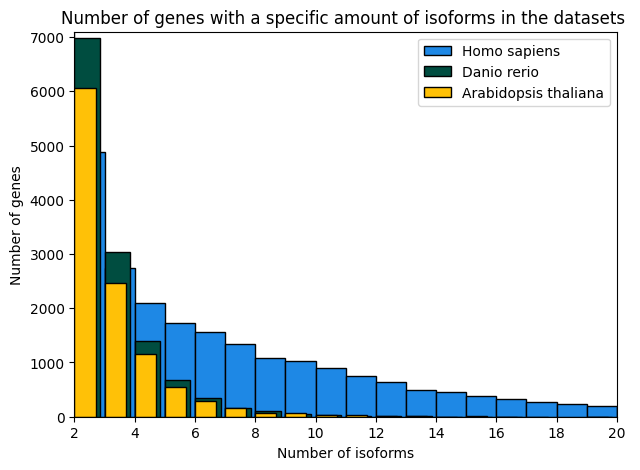

Figure S2: Number of genes with 2-20 isoforms. The number of genes with only one transcript was 35553 (61.6%) for the human reference, 23025 (67.8%) for the Arabidopsis reference and 16223 (55.8%) for the zebrafish reference. A maximum number of 82, 27 and 20 transcripts for one gene occurred in the human, Arabidopsis and zebrafish reference, respectively.

a) *Homo sapiens*

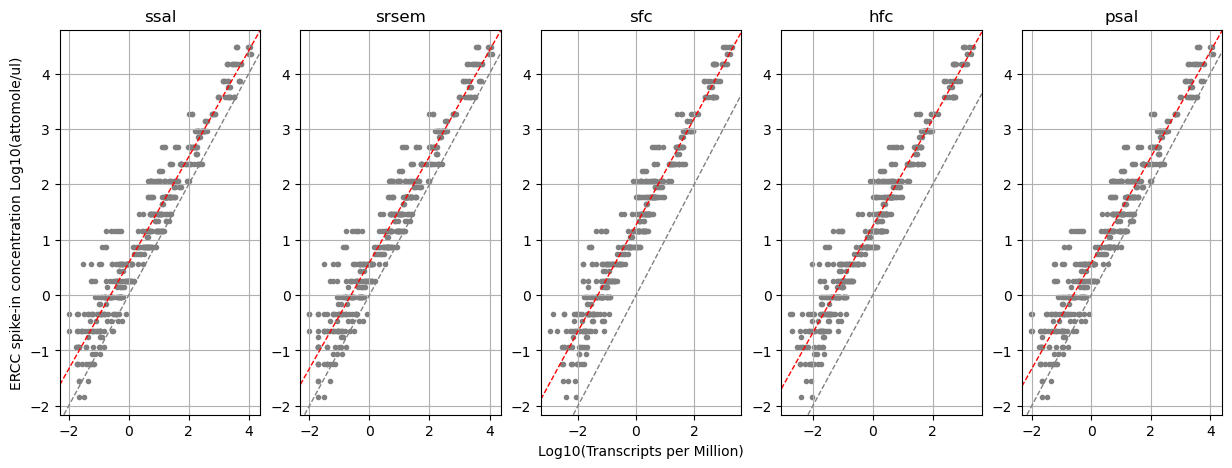

b) *Arabidopsis thaliana*

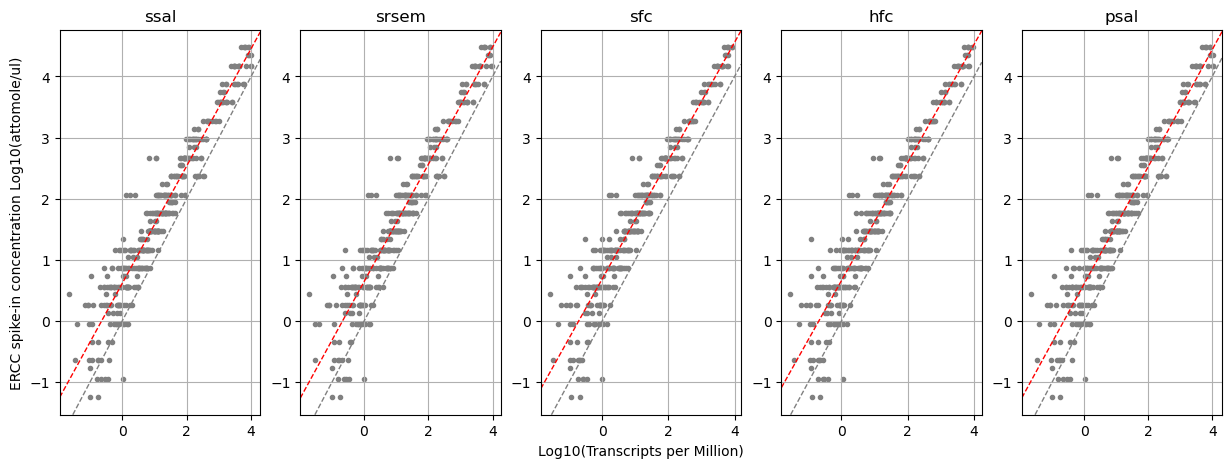

c) *Danio rerio*

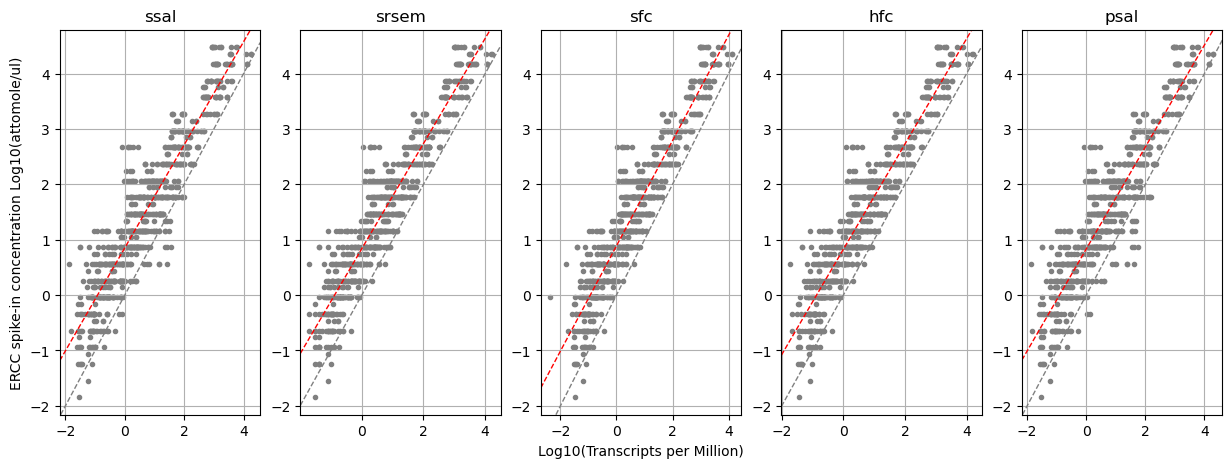

Figure S3: Visualization of dynamic range and lower limit of detection with ERCC RNA spike-ins for all datasets (a: human cell dataset, b: Arabidopsis dataset, c: zebrafish) and each pipeline setting (ssal: star+salmon v.3.2, srsem: star+rsem v.3.2, sfc: star+featurecounts v.1.4.2, hfc: hisat+featurecounts v.1.4.2, psal: pseudo-aligner salmon v.3.2). The red line represents the linear regression, the gray line indicates the 45° reference line. Values for slope and LLD are listed in Table 2.
